# Physiological System Dysregulation in Gene Expression Correlates Negatively with Age

**DOI:** 10.1101/2020.09.09.289389

**Authors:** Frédérik Dufour, Pierre-Étienne Jacques, Alan A. Cohen

**Author notes:** **For correspondence** (Alan A. Cohen).

## Abstract

We attempted to identify gene expression systems that dysregulate with age in human peripheral blood samples across five public datasets. Dysregulation of gene ontology (GO)-defined systems was measured using the Mahalanobis distance (D_M_), a measure of multivariant aberrance. We expected many weak positive D_M_-age correlations, indicating loss of homeostatic control. Out of the 5180 GO terms tested, we found 230 systems that replicated in at least three datasets. Surprisingly, all 230 systems showed negative D_M_-age correlations, in contrast to findings with clinical biomarkers. These systems were mostly metabolic functions related to small molecules and nitrogen compounds, transport functions, biosynthetic processes and response to stress functions. These results suggest a loss of responsiveness in these gene expression systems during the aging process, and contrast to some previous literature showing increased gene expression heterogeneity with age.

## Introduction

Aging can be characterized as a general loss of homeostasis in complex regulatory networks maintaining physiological functions (***Cohen, 2016***). This is accompanied by an accumulation of molecular damage and a progressive decline with age in cellular functions (***Kirkwood, 2005***). Many studies have looked at the change in expression variability during aging in different tissues and species, but there are conflicting conclusions on the subject. Some studies found a majority of genes that increase variance (***Işıldak et al., 2020***; ***Li et al., 2009***; ***Kedlian et al., 2019***), other studies found the variance mostly decreased with age (***Viñuela et al., 2018***), while other studies show there is an equal amount of genes that increase or decrease variance with aging (***Brinkmeyer-Langford et al., 2016***). However, few studies have looked at the change in variance during aging from a multivariate perspective. From a complex systems theory standpoint, the intricate interactions between the different genes create co-expression networks that can produce emergent properties of the systems that cannot be explained by the looking at the patterns of individual genes (***Cohen, 2016***; ***Kauffman et al., 1993***). Therefore, multivariate measures of variability could potentially bring further insights into the changes in variability during the aging process.

The Mahalanobis distance (D_M_) is a multivariate statistical distance that quantifies how aberrant a profile (e.g. of biomarkers) is compared to a reference population’s distribution, taking into account the correlation structure of the data (***Mahalanobis, 1936***). D_M_ has previously been used with transcriptomic data as a measure of gene expression aberrancy to study population differences (***Zeng et al., 2015***), gene expression functional properties (***Zeng et al., 2015***), brain aging (***Brinkmeyer-Langford et al., 2016***), breast cancer (***Schissler et al., 2015***), schizophrenia (***Huang et al., 2020***; ***Guan et al., 2019***), autism (***Guan et al., 2016***, ***2019***) and bipolar disease (***Guan et al., 2019***). D_M_ has also been extensively used with clinical blood biomarkers to study aging. It was shown to increase with age, suggesting that these systems dysregulate (lose homeostasis) during aging (***Li et al., 2015***). The dysregulation of these systems was associated with mortality, frailty, cardiovascular diseases and diabetes (***Milot et al., 2014***; ***Li et al., 2015***). One study has demonstrated that D_M_ measured globally and on different biomarker sets poorly correlated with one another, suggesting that the dysregulation of systems proceeds rather independently across systems (***Li et al., 2015***). Likewise, D_M_ and other biological age measures are also poorly related (***Belsky et al., 2017***). This suggests that the measures of biological age quantify different underlying processes during aging. Therefore, to have a global view of aging processes and more specifically dysregulation in humans, additional studies on dysregulation of systems during aging are needed, particularly at a finer biological scale.

In this study, we used D_M_ on microarray data from blood samples to look at the change in multivariate dispersion of gene expression with age. As with the blood biomarkers in previous studies, we hypothesized that we should find mostly positive associations between D_M_ and age, because of the loss of transcriptional control documented during the aging process. Few negative associations would be detected. We used five independent datasets and focused on results that replicate in multiple datasets, in order to avoid generating false positives or population-specific findings. Surprisingly, we found that correlations that were replicated in multiple datasets were exclusively negative, suggesting a loss of responsiveness in gene expression.

## Methods

### Datasets

The datasets used in this article were selected from the Gene Expression Omnibus (GEO) (***Clough and Barrett, 2016***) or ArrayExpress (***Sarkans et al., 2005***) databases. They were selected if they had at least 200 samples measured on the Illumina Expression BeadChips with complete age, sex and batch identification variables provided, after filtering out samples with known serious health conditions such as cancer or auto-immune diseases. In order to have appropriate age distributions, datasets that did not include individuals older than 75 years old or had narrow age distributions were also discarded. A total of five datasets listed in ***Table 1*** were therefore included in the analysis (***Beutner et al., 2011***; ***Sood et al., 2015***; ***Wingo and Gibson, 2015***; ***Peters et al., 2015***; ***Tarn et al., 2019***). Age ranges are large for all datasets, and one dataset (UKPSSR) is composed of almost exclusively female samples (***Appendix 1 Figure 1***). All datasets were measured on the version four of the Illumina HumanHT-12 Beadarray, except AddNeuro which also includes samples measured on the version three. For this dataset, the data were adjusted to remove the platform effects with the empirical Bayes adjustment algorithm, ComBat (***Johnson et al., 2007***), implemented in the sva package (version 3.34.0) (***Leek et al., 2019***).

**Table 1.** Dataset characteristics used in this study.

| **Dataset** | **Accession number** | **N samples included** | **N females (%)** | **Median age (min - max)** | **Region** |
| --- | --- | --- | --- | --- | --- |
| Leipzig LIFE Heart Study (Leipzig) | GSE65907 | 2112 | 725 (34.33) | 65 (25 - 97) | Leipzig, Germany |
| AddNeuroMed Consortium (AddNeuro) | GSE63063 | 714 | 431 (60.36) | 76 (52 - 100) | cross-European |
| Emory University Center for Health Discovery and Well-being (CHDWB) | GSE61672 | 332 | 222 (66.87) | 50 (25 - 78) | Atlanta, GA, US |
| Grady Trauma Project (GTP) | GSE58137 | 242 | 159 (65.7) | 45.2 (20 - 77) | Atlanta, GA, US |
| UK Primary Sjögren's Syndrome Registry (UKPSSR) | E-MTAB-8277 | 228 | 208 (91.23) | 57 (29 - 100) | United Kingdom |

### Methodological note

In the interest of transparency, we note that the description of the rest of this article, which follows standard conventions, ignores an important portion of the actual analyses conducted. We attempt to summarize them briefly here. First, we tested many different preprocessing methods in only one dataset (Rotterdam Study (***Ikram et al., 2017***), not included here). The results initially confirmed our hypothesis that D_M_ would increase with age in many systems; before publishing, we decided to replicate in an independent dataset (Leipzig). However, there was almost no overlap between the studies. This led to extensive exploration of how batch effects might bias the estimation of the correlation matrix, needed to calculate D_M_, and methods that might correct for batch effects. In the end, we concluded that there were no sufficient correction methods when batch is unknown, and limited ourselves to datasets with batches identified, thus excluding Rotterdam. We further identified that age structure could create biases and added a correction (i.e. adjusting the gene expression data for age).

### Data analyses

All analyses were run with R version 3.6.2 (***R Core Team, 2019***) and Bioconductor version 3.10 (***Reimers and Carey, 2006***). Analyses were replicated by an independent analyst to ensure programmatic and analytical reproducibility. Data transformations and figures were done using packages from the *tidyverse* (version 1.3.0) (***Wickham et al., 2019***) or the *corrplot* (version 0.84) (***Wei and Simko, 2017***) package.

### Data preprocessing

Before processing the data, hierarchical clustering analyses and visual inspection of the data distributions from randomly chosen probes, were done to remove any problematic samples in the datasets. Two individuals were removed in the UKPSSR dataset. Background subtraction was then applied on the raw data with the robust multi-array average (RMA) algorithm (*limma* package, version 3.42.2, (***Ritchie et al., 2015***)). Next, the data were transformed to log2 scale so that the distributions are approximately normal and adjusted for age and sex prior to batch normalization with the ComBat algorithm. It was important to adjust for age because we found D_M_ is sensitive to the age distribution of the datasets, which can bias the location of the reference population’s centroid (see Multivariate measure of system dysregulation) to the average age in the dataset (data not shown). Thus, D_M_ would change with age based on the age distribution if the data were not adjusted. Ideally, we would have uniform age distributions. Therefore, age and sex adjustments were done with *limma’s* lmFit function. Each probe was modeled independently as a function of age with spline models (splines package (version 3.6.3) bs function with three degrees of freedom) and sex as a covariate. For the batch normalization with ComBat, we used the residuals from these models as the input data. The beadchip plate identification numbers were used as batch identification for the Leipzig, AddNeuro and UKPSSR datasets, which means that batch size has a maximum of 12 samples per batch. For CHDWB, the “hybridization batch” covariate included four analysis batches of sizes 34, 59, 53 and 186 samples, respectively. For GTP, we created the “batch” variable by concatenating the amplification (“plateidamplification”) and hybridization (“plateidhybridization”) identification variables, for a total of 18 batches.

### Probe filtering

The *IlluminaHumanv4.db* package (version 1.26.0) (***Dunning et al., 2015***) was used to annotate the probes, estimate detection p-values and filter out bad quality probes. This package assigns a quality grade to each probe, based on the alignment of the Illumina probe sequences to the genome (***Barbosa-Morais et al., 2010***). Probes are scored with either a “Perfect”, “Good”, “Bad” or “No Match” quality grade. After processing the data, detection p-values were calculated with the calculateDetection function from the *beadarray* package (version 2.36.1) (***Dunning et al., 2007***). Given that some datasets did not include the control probes with the data, for all datasets we set controls to be probes with a quality score of “No match” to estimate the p-values. Probes that had no samples with a detection p-value below 0.05 were excluded from the analysis. Next, we filtered out the “Bad” and “No Match” quality graded probes as well as the probes hybridizing on sex chromosomes. Only probes available in all datasets after filtering were kept in the analysis.

### Gene ontology to define biological systems

We used gene ontology (GO) (***Ashburner et al., 2000***), more specifically the biological process (GO - BP) category of terms, to group probes into what we call functional systems. The GO index file (go-basic.obo) was downloaded from the Gene Ontology Consortium website in November 2019 (monthly release: 2019-10-07) and contains 47 375 terms. Probes were assigned to groups using the GO annotation file (GAF, monthly release 2019-10-07), also downloaded from the GO Consortium website at the same time. The probes mapped to a given gene were not always well-correlated and were therefore all included in a functional system. (Furthermore, D_M_ automatically eliminates redundancy among included variables.) The size of the groups was limited to a minimum of ten probes with at least two unique genes and a maximum of 228 probes (corresponding to the smallest dataset size, in order to be able to invert the covariance matrix and have all the same terms tested in all datasets). Overall, 5180 functional systems were tested (***Appendix 1 Figure 2***).

### Multivariate measure of system dysregulation

To evaluate the dysregulation of systems, we use the Mahalanobis distance (D_M_), a multivariate distance which measures the deviation of a biomarker profile from the reference population’s centroid, taking into account the correlation structure of the biomarkers in the system. D_M_ is calculated with the equation:

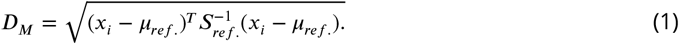

where *x_i_* is the vector of expression values in the functional system tested for individual *i*, *μ_ref._* is the vector of probe medians estimated from the reference population and 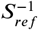 is the inverted reference population covariance matrix. For this study, we use each complete dataset as its own reference population. We conducted three sets of sensitivity analyses in which all datasets shared the same reference population: (1) all raw datasets combined and processed; (2) Leipzig (largest); (3) AddNeuro (2nd largest), which do not change the overall conclusions (***Figure 1***–***Figure Supplement 1***, ***Figure 1***–***Figure Supplement 2***, ***Figure 1***–***Figure Supplement 3***, ***Figure 2***–***Figure Supplement 1***, ***Figure 3***–***Figure Supplement 1***, ***Figure 3***–***Figure Supplement 2***, ***Figure 3***–***Figure Supplement 3***). D_M_ are then transformed to log scale, as they are approximately log-normally distributed.

### Assessing change in dysregulation with age

In order to identify systems that dysregulate with age using cross-sectional data, we ran correlation analyses between D_M_ and age for each system tested. Because we were initially interested in how many systems show significant associations, rather than which systems, p-value adjustments such as Benjamini-Hochberg are not appropriate. Rather, we compared the number of replicated correlations significant at *α* = 0.05 with the expected number of false positives (number of tests / 20). The overlap of results between datasets was tested using the exact test for multi-set intersections, implemented in the *SuperExactTest* package (version 1.0.7) (***Wang et al., 2019***). Results were considered to be overlapping if the correlations with age were significant with p < 0.05 and if they had the same direction.

### GO slim enrichment analyses

GO slims are higher-order categorizations of systems than the GO-BP used to calculate D_M_. We used GO slims to interpret the biological importance of systems that replicated across datasets. The GO slims were obtained from the goslim-generic.obo file (version 1.2), downloaded from the GO Consortium website in March 2020. GO terms that overlap in at least three datasets were mapped to the GO slims and counted. If a term mapped to more than one GO slim, all the correlations were kept and counted. The mapping to GO slims was also conducted on all the GO systems tested, to determine the expected proportions of GO terms mapping to the GO slims. One-tailed z-tests of proportions, with no Yates continuity corrections were done to compare the observed and expected proportions. P-values were adjusted for multiple comparisons with the Benjamini-Hochberg method.

## Results

### D_M_ correlations with age within datasets

After processing the data and filtering out probes, a total of 21 591 probes (~45%) were retained and grouped into functional systems using GO as described in Methods. Correlation analyses between D_M_ and age were conducted on a total of 5180 systems. ***Figure 1*** shows the number of correlations significant at *α* = 0.05. For all datasets, the vast majority of the significant correlations with age are negative, and above the number expected by chance (except for UKPSSR). In contrast, the number of positive age correlations is less than what is expected by chance for all datasets. Qualitatively, these results only change minimally when the reference population is changed (***Figure 1***–***Figure Supplement 1***, ***Figure 1***–***Figure Supplement 2***, ***Figure 1***–***Figure Supplement 3***).

**Figure 1.**
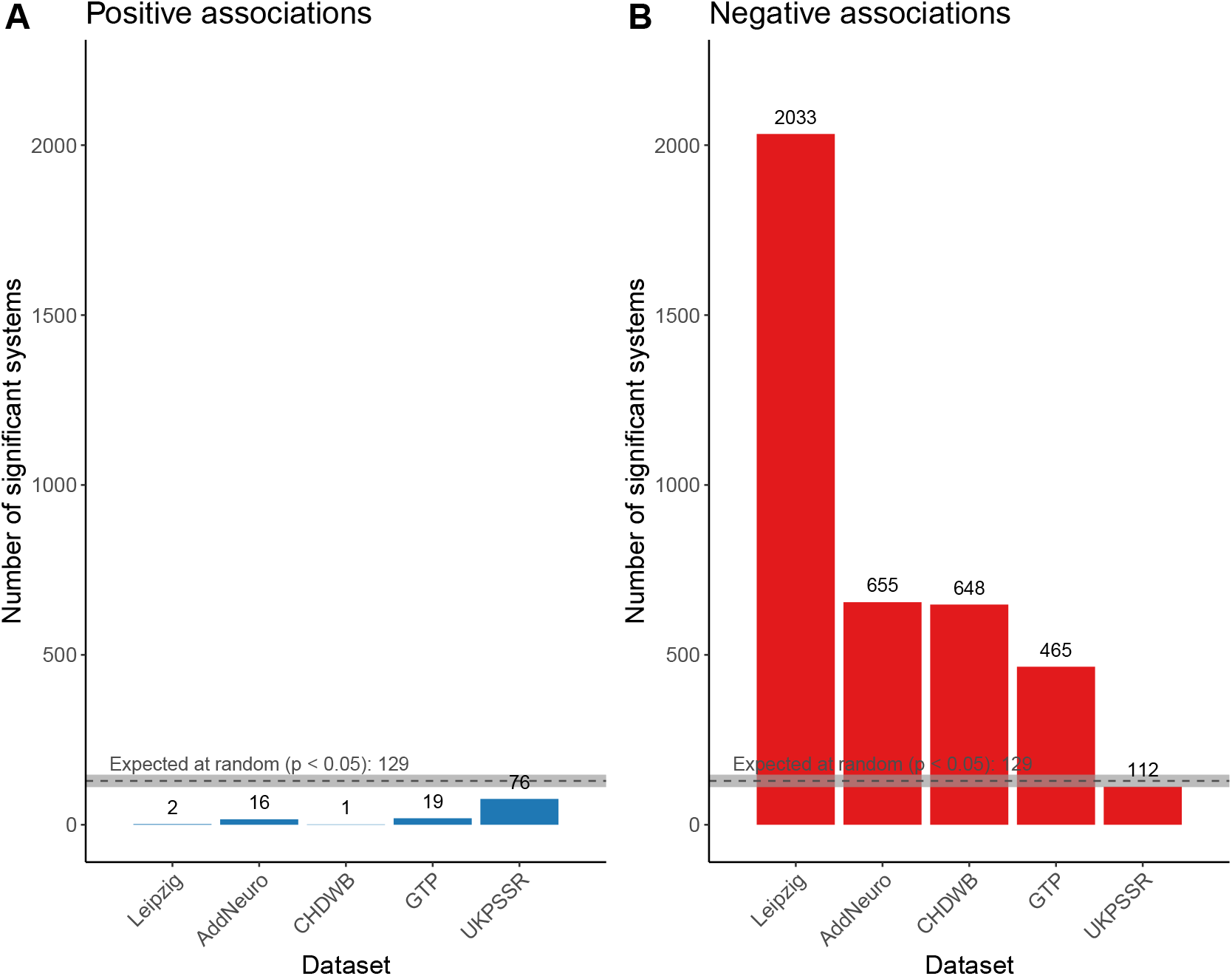
D_M_ tends to decrease with age across the systems tested. The graph tallies the number of significant correlations (*α* = 0.05) for each dataset. The positive and negative correlations are in blue and red, respectively. Given the 5180 systems tested, we should expect 129 significant correlations (grey dotted line), by chance alone and with equal chances of having positive or negative correlations. This estimation is based on 1000 random samples of 5180 p-values from the uniform distribution U(0,1), from which we counted the number of p-values < 0.025 (0.05 (*α* threshold) X 0.5 (positive vs. negative correlations)). The grey area around the dotted line is the 95 % confidence interval for the null distribution. **Figure 1–Figure supplement 1.** Number of significant correlations when the reference population is all datasets combined. **Figure 1–Figure supplement 2.** Number of significant correlations when the reference population is the Leipzig dataset for all other datasets. **Figure 1–Figure supplement 3.** Number of significant correlations when the reference population is the AddNeuro dataset for all other datasets.

### Overlap between datasets

Next, we tested the overlap across datasets of the significant correlations, for all possible combinations of the five datasets. Here, we consider that a functional system overlaps between two datasets when the correlation signs are in the same direction and the relationship is significant in both datasets. In total, 230 unique systems were replicated in at least three datasets and were considered to be potentially interesting systems associated with aging (Supplementary File 1). Interestingly, all the 230 correlation coefficients are negative. We find five systems that are replicated in all five datasets (p-value = 1.21e-7, ***Table 2***). For the other combinations of the five datasets, we find the overlap is mostly significant, even after adjusting the p-values for multiple comparisons (***Figure 2***). When the reference population is changed, the replicated systems all remain negatively correlated with age (Supplementary File 1). However, the identities of the replicated systems tend to vary (***Figure 2***–***Figure Supplement 1***). We also calculated Spearman rank correlations between the correlation coefficients of dataset pairs, showing poor agreement in changes of D_M_ with age over the 5180 systems (***Figure 2***–***Figure Supplement 2***).

**Table 2.** Functional systems replicated in all 1ve datasets.

| GO | Term | Group size<br>(unique genes) | Correlation sign |
| --- | --- | --- | --- |
| GO:0032496 | response to<br>lipopolysaccharide | 201 (139) | Negative |
| GO:0044262 | cellular carbohydrate<br>metabolic process | 197 (124) | Negative |
| GO:0050663 | cytokine secretion | 228 (140) | Negative |
| GO:0050707 | regulation of cytokine<br>secretion | 205 (126) | Negative |
| GO:0072593 | reactive oxygen<br>species metabolic<br>process | 218 (139) | Negative |

**Figure 2.**
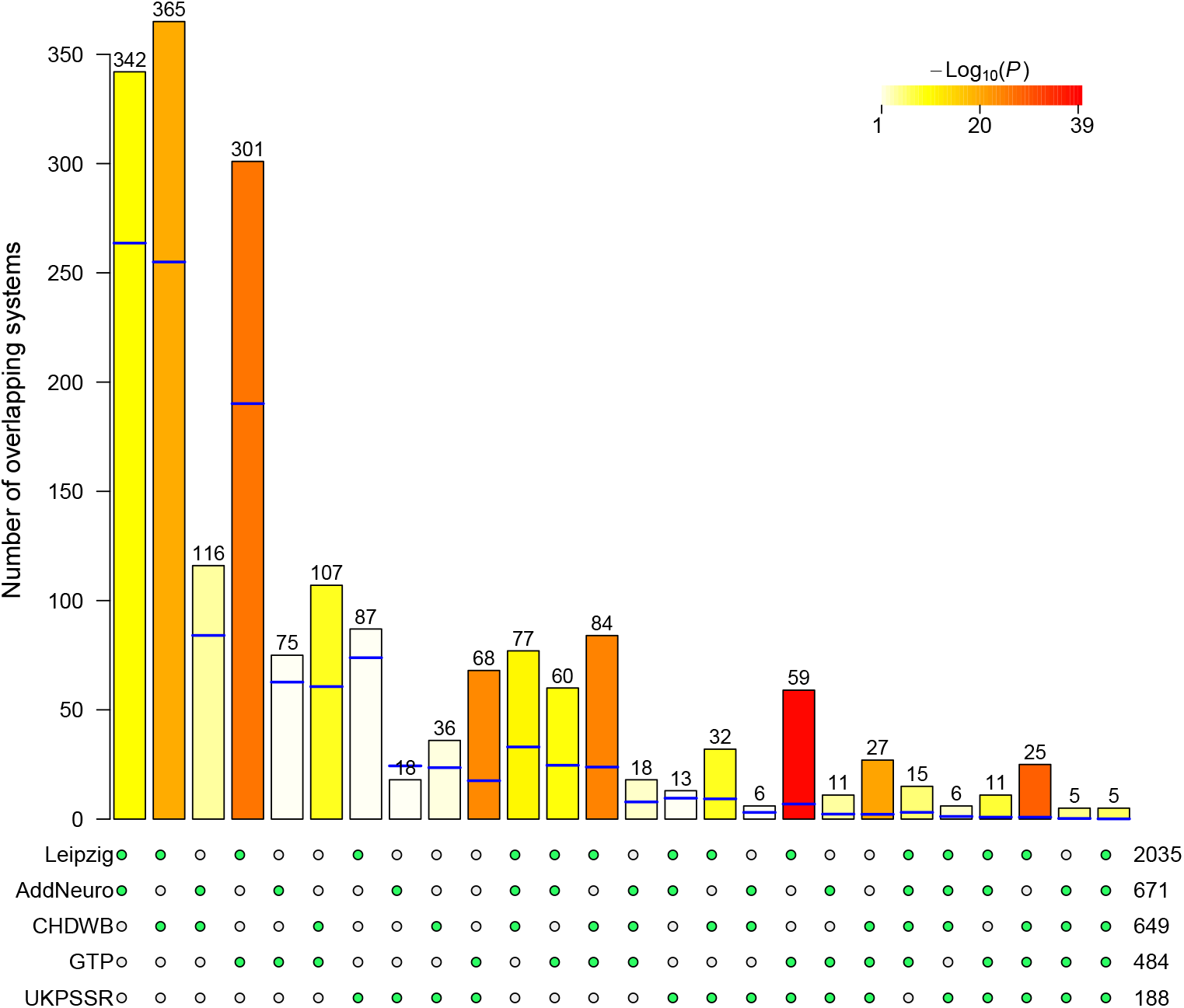
The significant correlations that replicate across datasets. The number of age-associated systems overlapping between datasets are represented by the bars in the graph. The blue lines represent the overlap expected by chance alone, given the number of systems tested (5180) and the number of significant correlations in the sets (shown to the right of the dot matrix). The color intensity is the negative log of the p-values from Fisher exact tests between expected and observed intersections. The color is white if the p-value > 0.05, after Benjamini-Hochberg adjustment. At the bottom of the graph, the matrix of green (included) and white (excluded) dots indicate the sets in the intersection. Only intersections which are not empty are shown in the barplot. **Figure 2–Figure supplement 1.** Overlap of replicated systems when changing the reference population. **Figure 2–Figure supplement 2.** Spearman rank correlation plot between D_M_-age correlation coefficients.

### Gene ontology of replicated systems

In order to get a more summarized view of the 230 replicated systems, we mapped them to GO slims (***Figure 3***). The replicated systems were mapped to 47 GO slims, substantially fewer than the 67 observed among the 5180 systems tested. The top five most represented GO slims are ‘cellular nitrogen compound metabolic process’ (34 terms), ‘transport’ (29 terms), ‘response to stress’ (20 terms), ‘small molecule metabolic process’ (18 terms) and ‘biosynthetic process’ (18 terms). Based on z-tests of proportions, ‘cellular nitrogen compound metabolic process’, ‘response to stress’, ‘chromosome organization’ and ‘vacuolar transport’ GO slim term are enriched compared to expected proportions. In contrast, ‘anatomical structure development’ associated systems are underrepresented among the 230 replicated systems. We find this more general profile of the replicated systems is similar when the reference population is varied (***Figure 3***–***Figure Supplement 1***, ***Figure 3***–***Figure Supplement 2***, ***Figure 3***–***Figure Supplement 3***).

**Figure 3.**
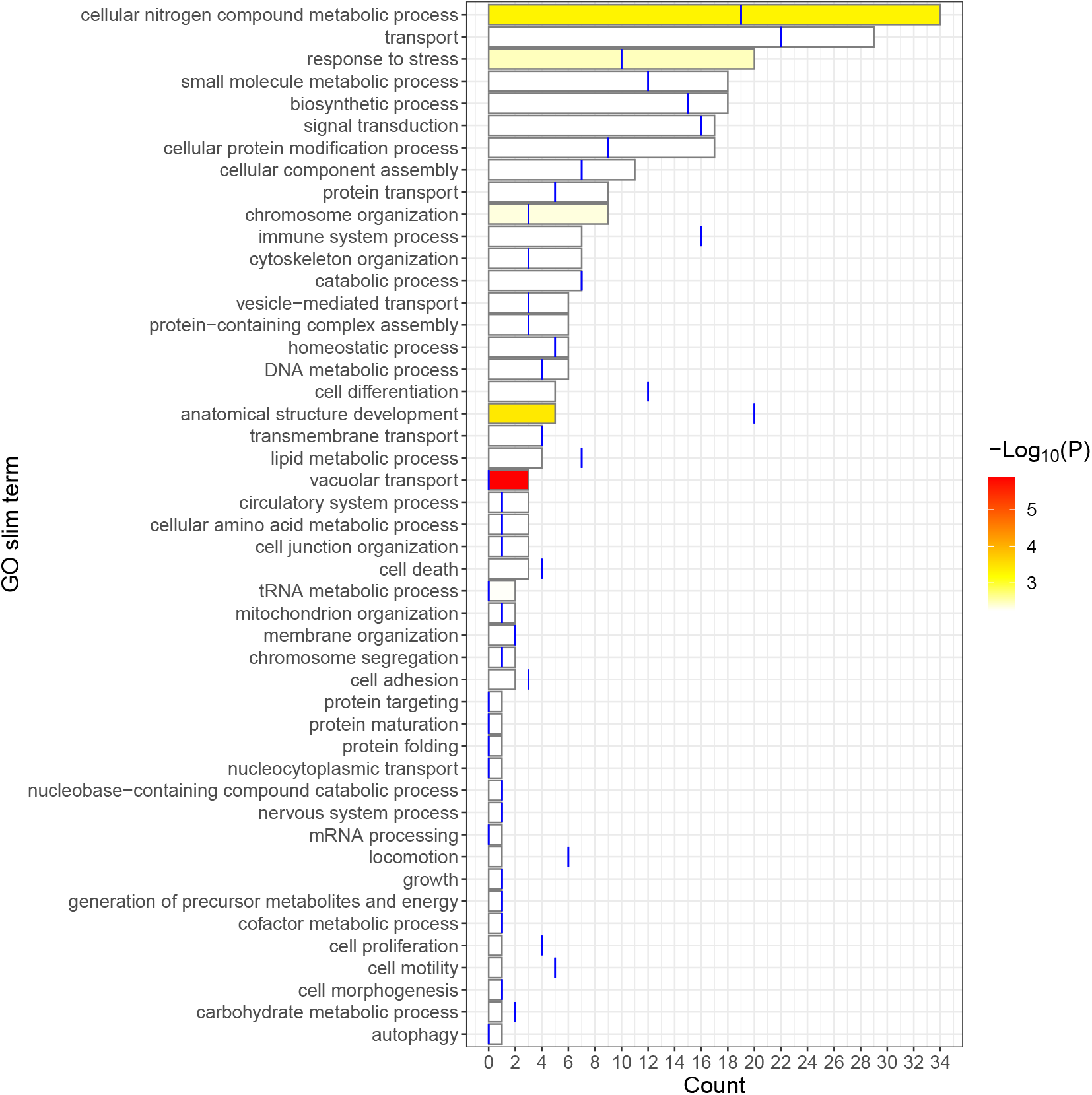
GO slim map and enrichment. The bar graph shows the number of GO terms, out of the 230 terms that overlap in three datasets, that are mapped to the GO slims listed. The expected number of systems mapping to the GO slims is given by the blue line. This was calculated by multiplying the expected proportions of the GO terms tested mapping to the GO slim with the number of unique systems overlapping in at least three datasets (230). The color intensity is the negative log of the p-values from the one-tailed z-tests of proportions. Color is set to white when p-values > 0.05 after adjusting for multiple comparisons with the Benjamini-Hochberg procedure. **Figure 3–Figure supplement 1.** GO slim map and enrichment when the reference population is all datasets combined. **Figure 3–Figure supplement 2.** GO slim map and enrichment when the reference population is the Leipzig dataset for all other datasets. **Figure 3–Figure supplement 3.** GO slim map and enrichment when the reference population is the AddNeuro dataset for all other datasets.

## Discussion

### Aging is associated with a general decrease in expression variability

In this study, we used a multivariate approach, more specifically D_M_, to look at integrative, system-level changes in gene expression variance during aging. Based on previous work with blood biomarkers, we expected a mix of mostly positive and a few negative associations between D_M_ and age. Systems with different functions need to be more or less responsive to changes in the internal and environmental conditions. Indeed, some systems, such as stress response systems or the immune system need to be highly responsive to changes in the environment (***Alemu et al., 2014***). Thus, during the aging process, we could expect a decrease in the association of D_M_ with age for these systems, which might indicate malfunction and a loss of responsiveness in the system. On the other hand, regulatory housekeeping functions probably need to be less variable (***Alemu et al., 2014***). Thus, an increase in D_M_ with age in these systems would suggest a loss in control over these systems. In our study, we found a global decreasing trend in D_M_ with age for all systems that were replicated in at least three datasets. Therefore, it seems that at the gene expression level, our method only captures the replicated negative correlations, though we cannot exclude that some positive correlations might still be present, but not detected/replicated. Nevertheless, it is also possible that, contrary to the blood biomarker level, gene expression systems only lose variability with age, when we use multivariate methods to assess variability.

There were a fair number of systems replicated in at least three datasets (n=230), more than in the study published by (***Brinkmeyer-Langford et al., 2016***) which found differentially variable systems (GO) associated with age in brain samples. In that study, they used very similar methods to ours and found five systems that showed decreases in variance and three that showed increases in variance. Out of these systems, one system (‘GO:0032214: negative regulation of telomere maintenance via semi-conservative replication’) is a child term of three of the systems replicated in our study (‘GO:0000723: telomere maintenance’, ‘GO:0032200: telomere organization’ and ‘GO:0032205: negative regulation of telomere maintenance’). These differences are most possibly explained by different tissues included in the analyses and the differences in the processing of the data prior to the analysis.

### The 230 age-associated systems represent many hallmarks of aging

The 230 age-associated systems represent very diverse cellular processes. Not surprisingly, we found some functions related to the regulation of blood circulation and heart contraction. We hypothesize that the loss of responsiveness in these systems could serve as a basis for understanding why heart rate variability slows with aging (***Stein et al., 2009***), although independent validation is needed. Many processes were related to cellular structure maintenance, organization and regulation, such as actin filaments, microtubules, adhesion and junctions, vesicle and vacuole, along with multiple processes related to vacuolar transport (ex. endocytosis, exocytosis, autophagy, phagocytosis, secretion and lysosomal functions). These are key components for the proper functioning of many immune cells. Furthermore, some GO terms relate specifically to the immune system, such as the regulation of innate and adaptive immune response, T cell activation, the production of interferon *β*, cytokine secretion and the response to protein from bacterial origin. The implication of the immune system is unsurprising in peripheral blood samples. Among the age-associated terms, many other stress response functions are represented. They include functions such as the response to oxidative stress, the response to decreasing oxygen levels, the regulation of superoxide metabolism, double strand break repair and the stress activated signaling pathways MAPK, ERAD, JNK, TGF*β*, TNF, NF-kappaB. A decrease in responsiveness in these systems could have dramatic effects on the organism, leaving it less robust to environmental and internal stressors.

Interestingly, among the age-associated systems, we find other systems associated with hallmarks of aging (***López-Otín et al., 2013***). Since we are looking at transcriptomic data, it is not surprising to find many DNA and RNA biosynthetic processes. We also found some RNA maturation (splicing) and processing functions, along with many functions related to the organization of the nucleus. Of particular interest, well documented hallmarks of aging, such as telomere maintenance functions, epigenetics and histone modifications, DNA packaging functions and protein-DNA complex organization are some of the processes we found were negatively associated with age. Another group of GO terms relate to the maintenance of proteostasis. They include the maturation, modification and localization of proteins in cells along with the removal of misfolded proteins (i.e. protein polyubiquitination, cellular response to topologically incorrect protein). One hallmark of aging that is well represented is mitochondria and energy production dysregulation. Terms related to this hallmark include mitochondria organization functions, the respiratory chain assembly and the metabolism reactive oxygen species. Finally, there are also some terms related to nutrient sensing dysfunctions, such as response to starvation, lipid metabolism, carbohydrate metabolism and response to nutrient levels.

### Limits of the study

Although there was a fair amount of overlap between the datasets, we did not find high correlations between the associations of D_M_ with age (***Figure 2***–***Figure Supplement 2***). This suggests that for most of the systems the signal might be hidden by substantial noise, making the results somewhat variable across the datasets. The UKPSSR dataset is the only dataset for which we did not find more significant negative correlations than expected by chance. This dataset contains primary Sjögren’s Syndrome patients, which is a common autoimmune rheumatic disease (***Fox, 2005***). It was not filtered out during the dataset selection process, but it is a clear demonstration of how the population composition could have confounding effects on the results. Other confounding factors such as chronic diseases or smoking are sometimes not specified in these cohorts, since, as with most secondary-omics analysis, we have very limited data on the samples. Therefore, there are very few sensitivity analyses that can be done. The choice of the reference population to estimate D_M_ seems to influence the identities of the replicated systems (***Figure 2***–***Figure Supplement 1***). However, our set of sensitivity analyses showed the results were qualitatively replicated when we changed the reference population. We believe this is likely a reflection of a larger set of systems that show true changes with age, but with the identity of the specific systems variable across analyses due to stochasticity and substantial noise in quantification and processing steps. Cell proportions, which vary during aging (***Valiathan et al., 2016***), is another non-negligible confounder. Indeed, different cell types have different expression profiles (***Abbas et al., 2005***) and different aging trajectories of variability (***Kimmel et al., 2019***). As with most cohort studies on aging, there is a possible survival bias in our results, where the individuals that survive into old ages are likely healthier than an average individual that did not survive to these ages. This would have the effect of underestimating the magnitude of changes in D_M_ with age.

It is difficult to fully dissect the microarray signals into technical noise and true biological signal. Many authors have attempted to compare different processing methods to find differentially expressed genes in different conditions (***Chen et al., 2011***; ***Müller et al., 2016***; ***Freytag et al., 2015***). However, in our work, we use the absolute expression signal as the data points. Thus, there might be some residual artifacts from the data processing steps. Here, we used ComBat to remove batch effects with small batch sizes (not greater than 12 samples per batch in some datasets). Based on simple simulations, by chance alone, this can introduce correlations between age and batches, confounding the effects of age on gene expression variability with batch effects (data not shown). After processing the data, we suppose that the estimates of the mean, variances and covariances obtained from cross-sectional studies are useful approximations of the within-individual changes with age. However, we do not formally test this hypothesis. To fully assess this limit, longitudinal data should be further explored.

To summarize, our analyses suggest many systems might lose responsiveness with increasing age. These systems are over-represented in biological functions known to change during aging, and represent a wide range of cellular processes that could leave the organism more vulnerable to physiological changes or environmental stressors. The balance between loss of responsiveness as implied here, and between loss of control, detected in many other studies, remains to be better understood.

## Code and data availability

R scripts used for the analyses of the data are available on our group’s Github page at https://github.com/cohenaginglab/Gene_expression_variability. Datasets were downloaded from the GEO (accession numbers: GSE65907, GSE63063, GSE61672, GSE58137) and ArrayExpress (accession numbers: E-MTAB-8277) databases.

## Acknowledgements

We would like to thank Francis. B. Lavoie that replicated the results to ensure programmatic and analytical reproducibility. We also thank the anonymous reviewers for their helpful feedback on earlier versions of the manuscript.

## Contributions

F.D., P.E.J. and A.A.C. conceptualized and developed the study. F.D. conducted the analyses, wrote the manuscript, and all authors contributed to revisions. This work is supported by Canadian Institutes of Health Research (CIHR) grant #153011, and National Science and Engineering Research Council Grant #RGPIN-2018-06096. A.A.C. has a Senior Fellowship and P.E.J. a Junior2 Fellowship from the Fonds de recherche du Québec – Santé (FRQ-S), and they both are members of the FRQ-S funded Centre de recherche du CHUS and Centre de recherche sur le vieillissement.

## Appendix 1

**Appendix 1 Figure 1.**
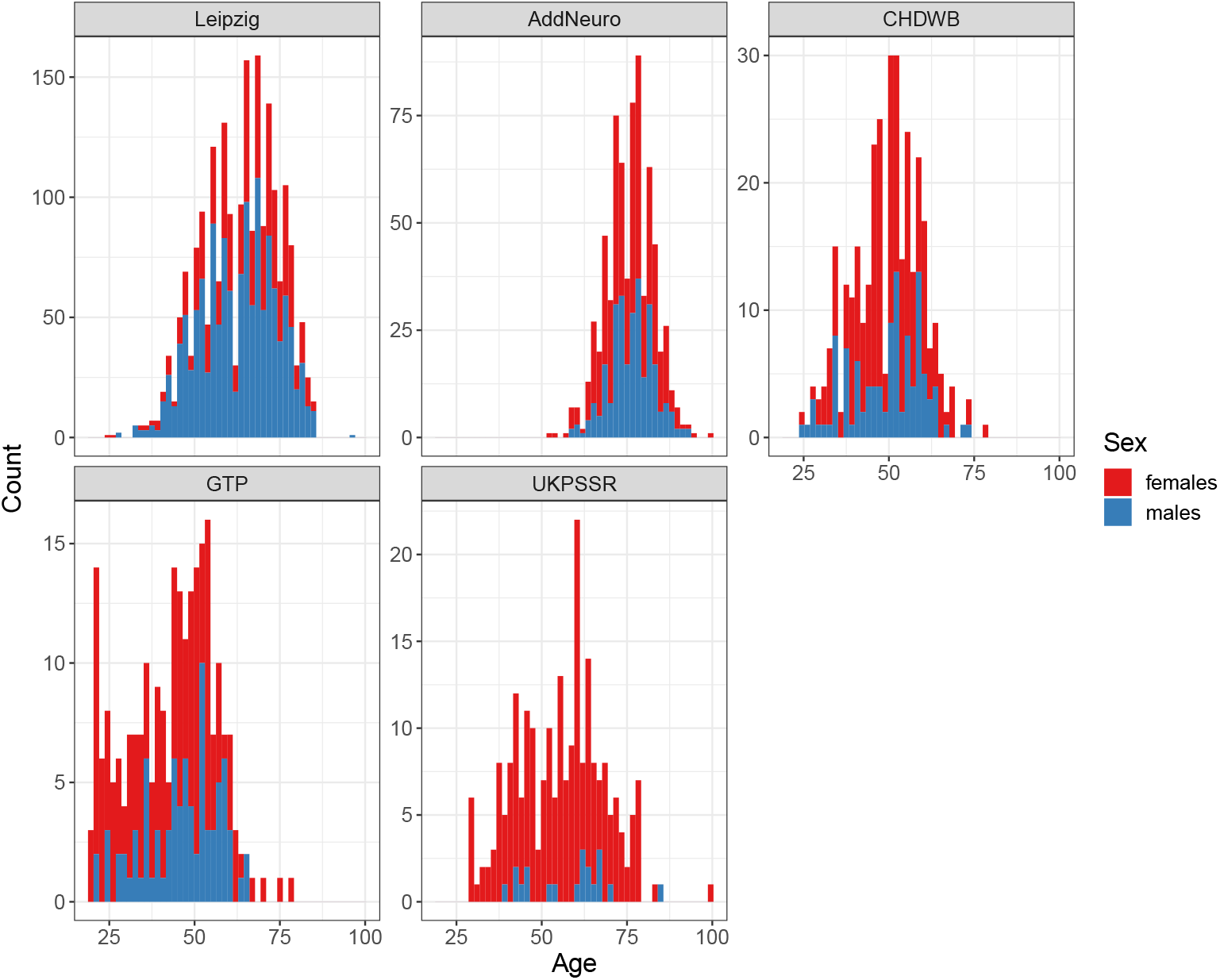
Age and sex distributions of the datasets. The bar graphs show the age and sex distributions in the different datasets. Colors give the proportions of females and males in red and blue, respectively.

**Appendix 1 Figure 2.**
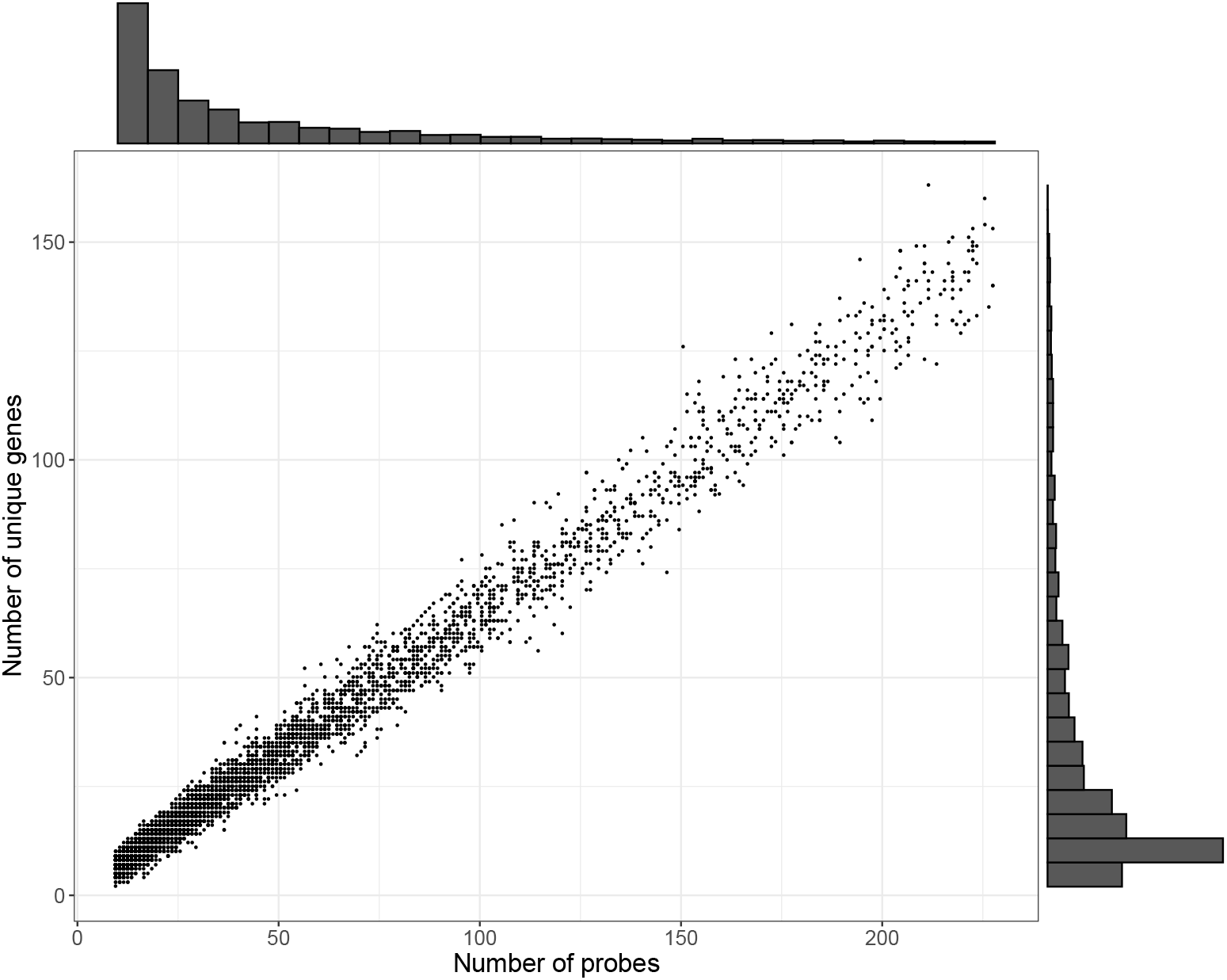
Group size distributions for the 5180 functional systems tested. The histograms show the group size distribution, based on the number of probes (top) and the number of unique genes (right) for the 5180 functional systems tested. The scatterplot shows the number of unique genes in each system.

---

**Figure 1–Figure supplement 1.**
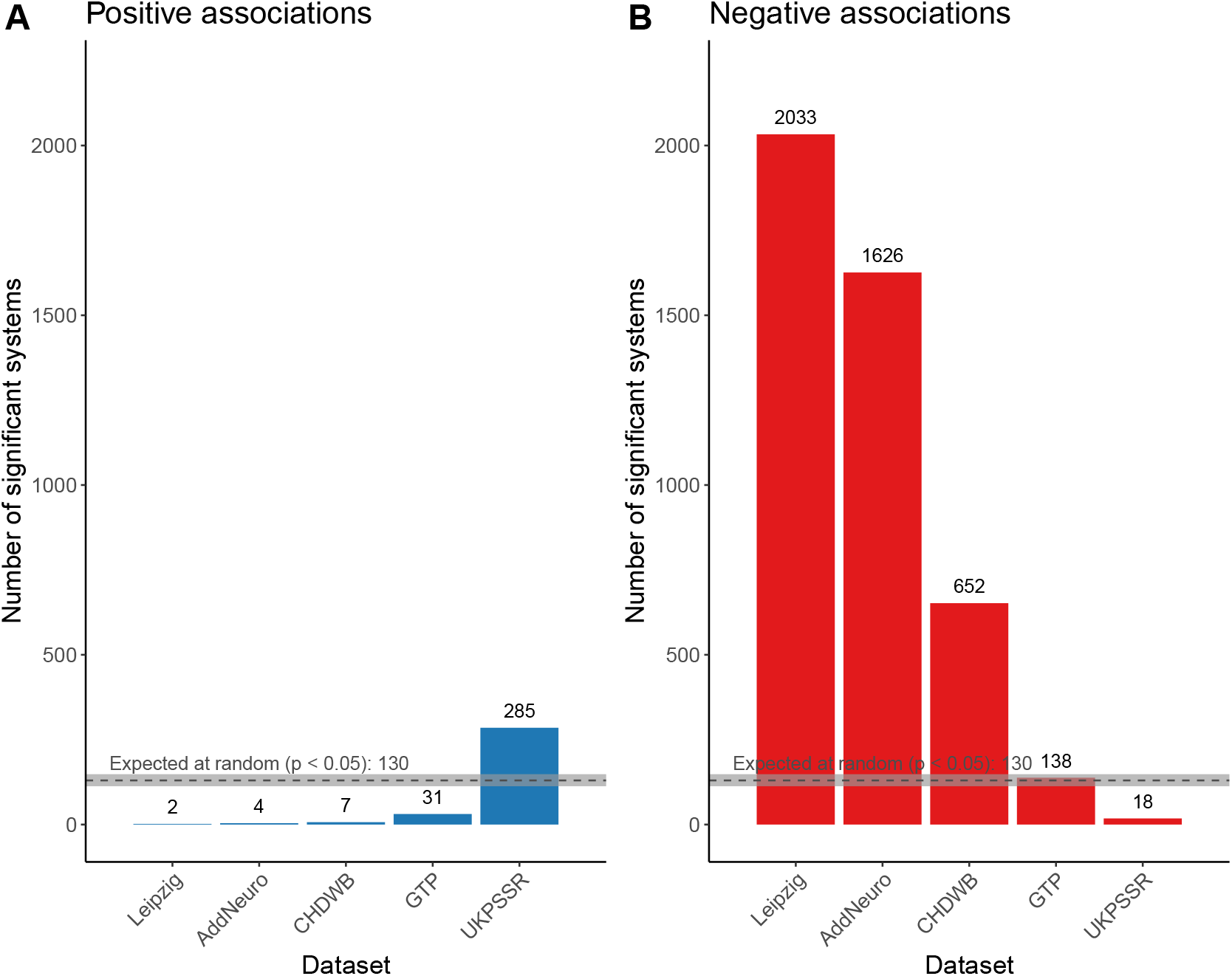
Number of significant correlations when the reference population is all datasets combined. The graph counts the number of significant correlations (*α* = 0.05) for each dataset, when the reference population is all datasets combined. The positive and negative correlations are in blue and red, respectively. Given the 5180 systems tested, we should expect 129 significant correlations (grey dotted line), by chance alone and with equal chances of having positive or negative correlations. This estimation is based on 1000 random samples of 5180 p-values from the uniform distribution U(0,1), from which we counted the number of p-values < 0.025 (0.05 (*α* threshold) X 0.5 (positive vs. negative correlations)). The grey area around the dotted line is the 95 % confidence interval for the null distribution.

**Figure 1–Figure supplement 2.**
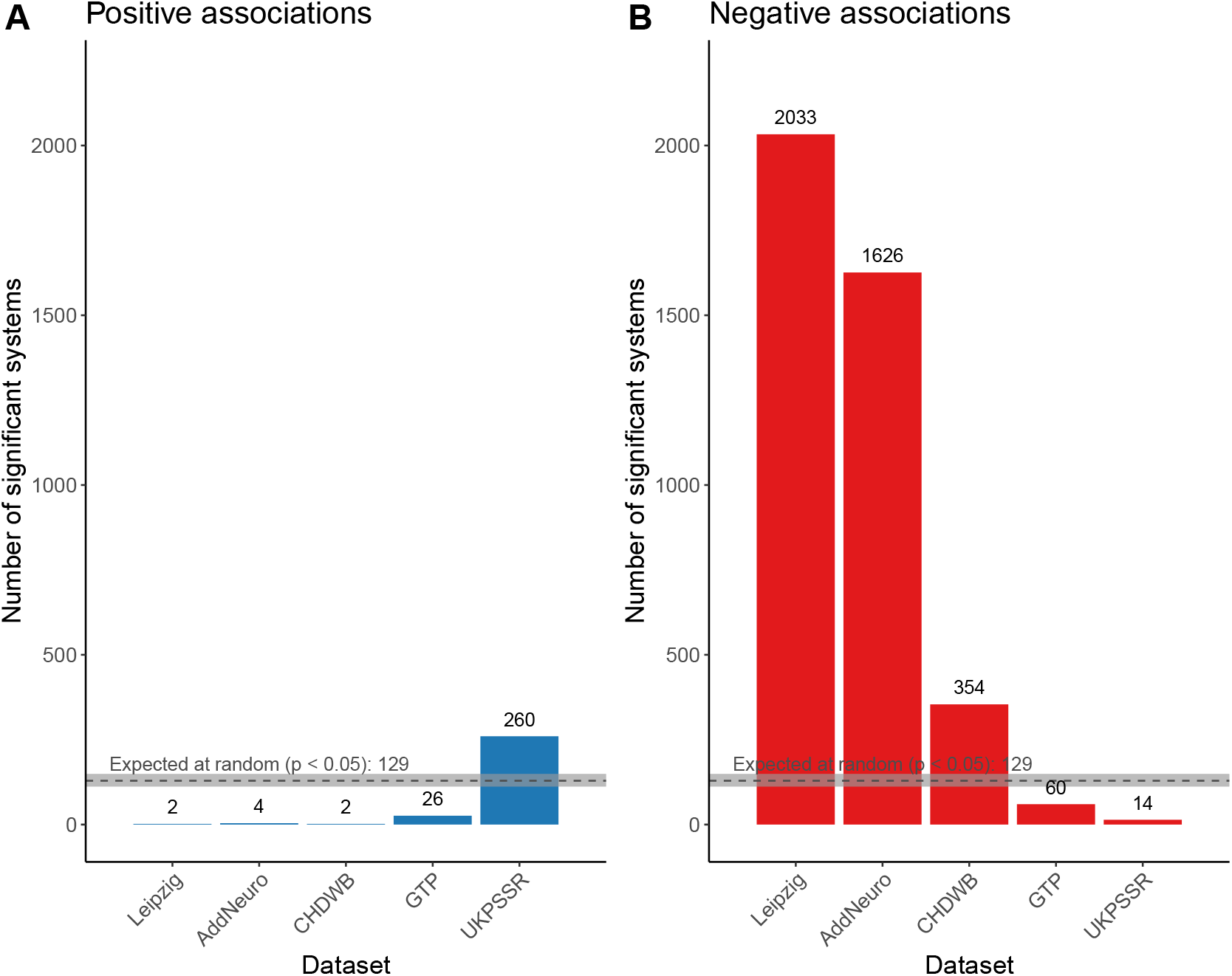
Number of significant correlations when the reference population is the Leipzig dataset for all other datasetss. The graph counts the number of significant correlations (*α* = 0.05) for each dataset, when the Leipzig dataset is the reference for all other datasets. The positive and negative correlations are in blue and red, respectively. Given the 5180 systems tested, we should expect 129 significant correlations (grey dotted line), by chance alone and with equal chances of having positive or negative correlations. This estimation is based on 1000 random samples of 5180 p-values from the uniform distribution U(0,1), from which we counted the number of p-values < 0.025 (0.05 (*α* threshold) X 0.5 (positive vs. negative correlations)). The grey area around the dotted line is the 95 % confidence interval for the null distribution.

**Figure 1–Figure supplement 3.**
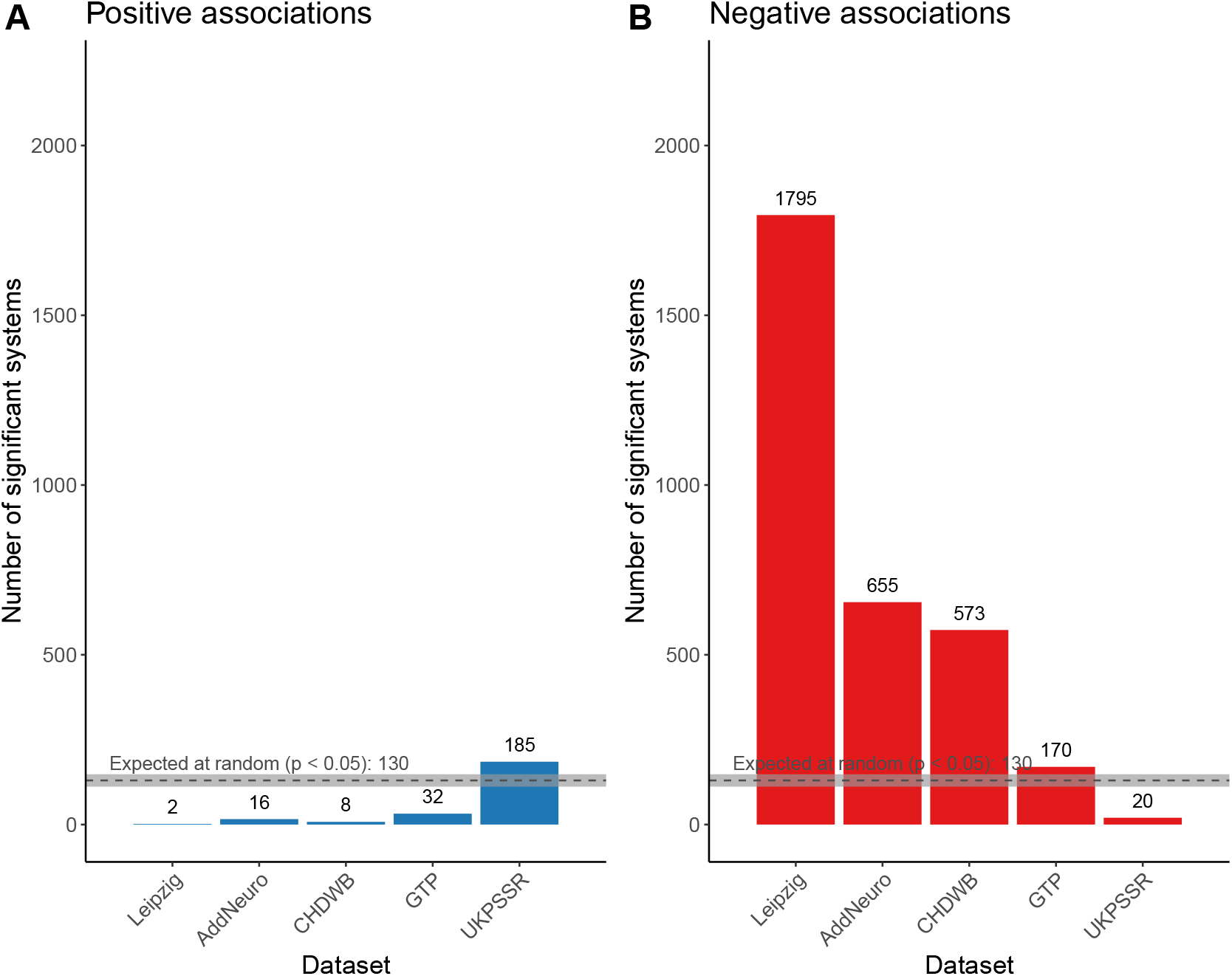
Number of significant correlations when the reference population is the AddNeuro dataset for all other datasets. The graph counts the number of significant correlations (*α* = 0.05) for each dataset, when the AddNeuro dataset is the reference for all other datasets. The positive and negative correlations are in blue and red, respectively. Given the 5180 systems tested, we should expect 129 significant correlations (grey dotted line), by chance alone and with equal chances of having positive or negative correlations. This estimation is based on 1000 random samples of 5180 p-values from the uniform distribution U(0,1), from which we counted the number of p-values < 0.025 (0.05 (*α* threshold) X 0.5 (positive vs. negative correlations)). The grey area around the dotted line is the 95 % confidence interval for the null distribution.

**Figure 2–Figure supplement 1.**
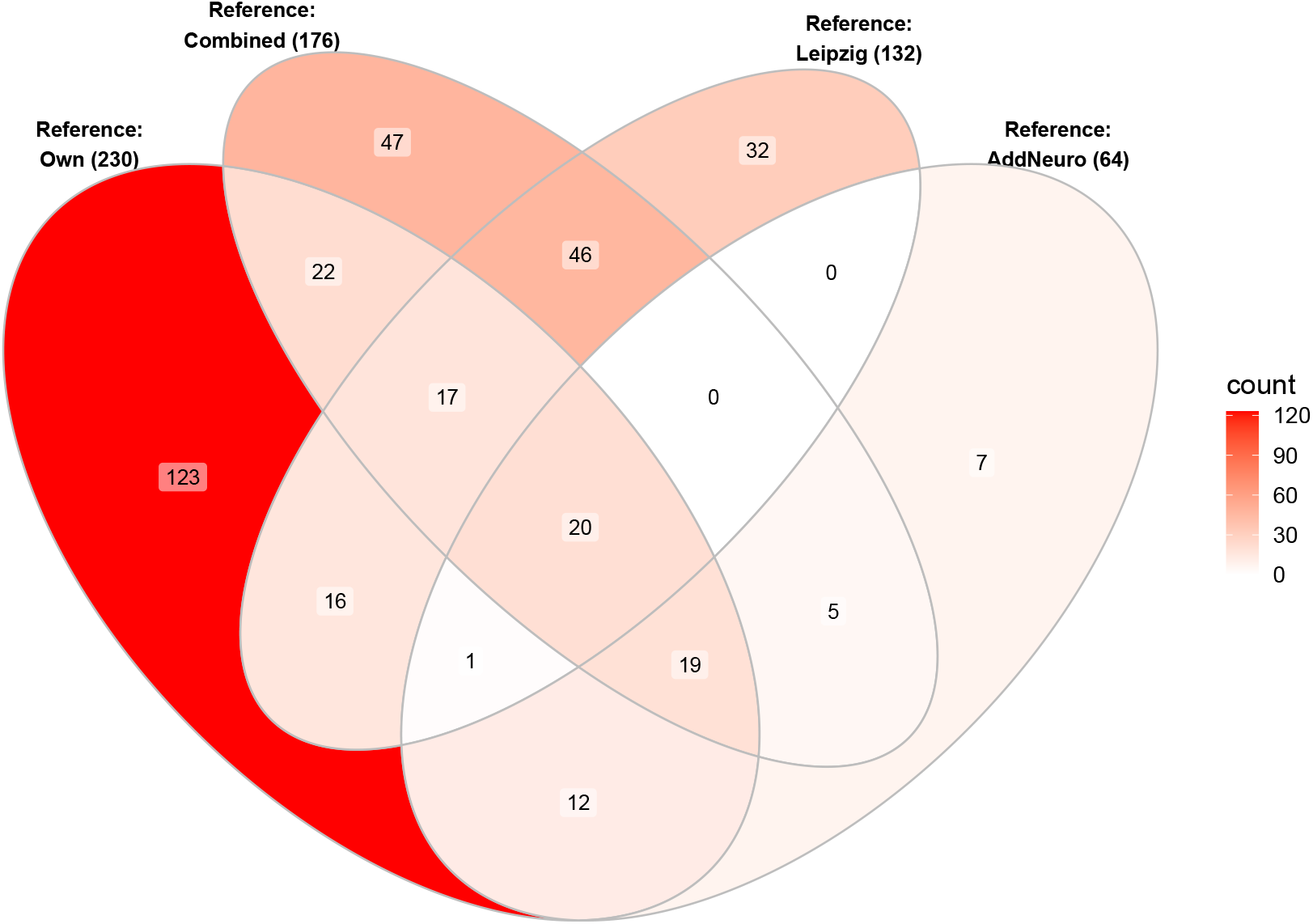
Overlap of replicated systems when changing the reference population. The venn diagram displays the intersection of replicated systems in at least three datasets when changing the reference population. The color scale is the count for the intersection. The number of systems replicated in at least 3 datasets in each set of sensitvity analyses is given by the number between parentheses in the labels.

**Figure 2–Figure supplement 2.**
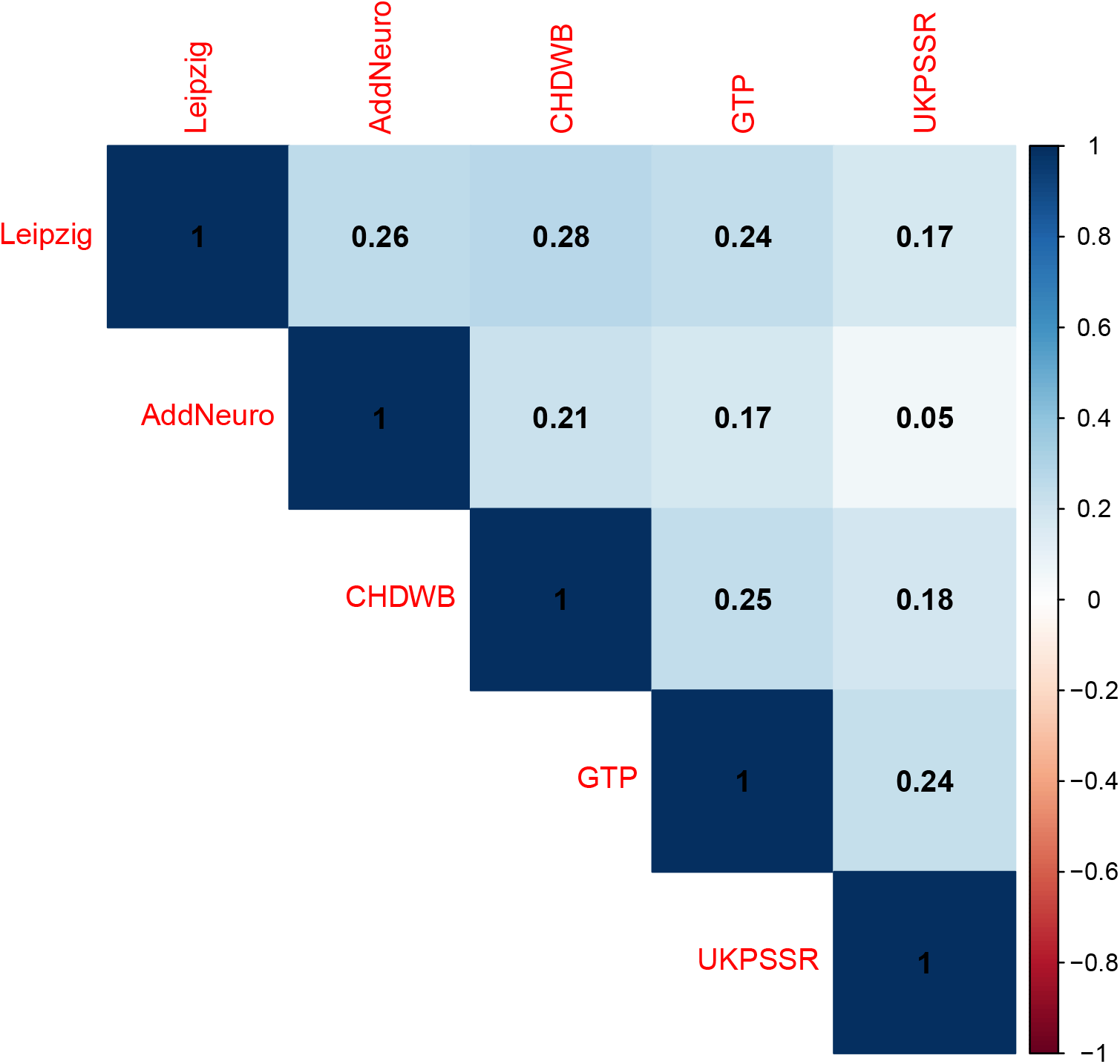
Spearman rank correlation plot between D_M_-age correlation coefficients. The correlation plot compares the relative overlap between datasets by looking at the similarity of the ranks across the 5180 systems tested. The numbers are the estimated *ρ* (rho) values and they are represented by the color intensities, blue and red being respectively positive and negative correlations. Non-significant correlations are indicated by an X in the square.

**Figure 3–Figure supplement 1.**
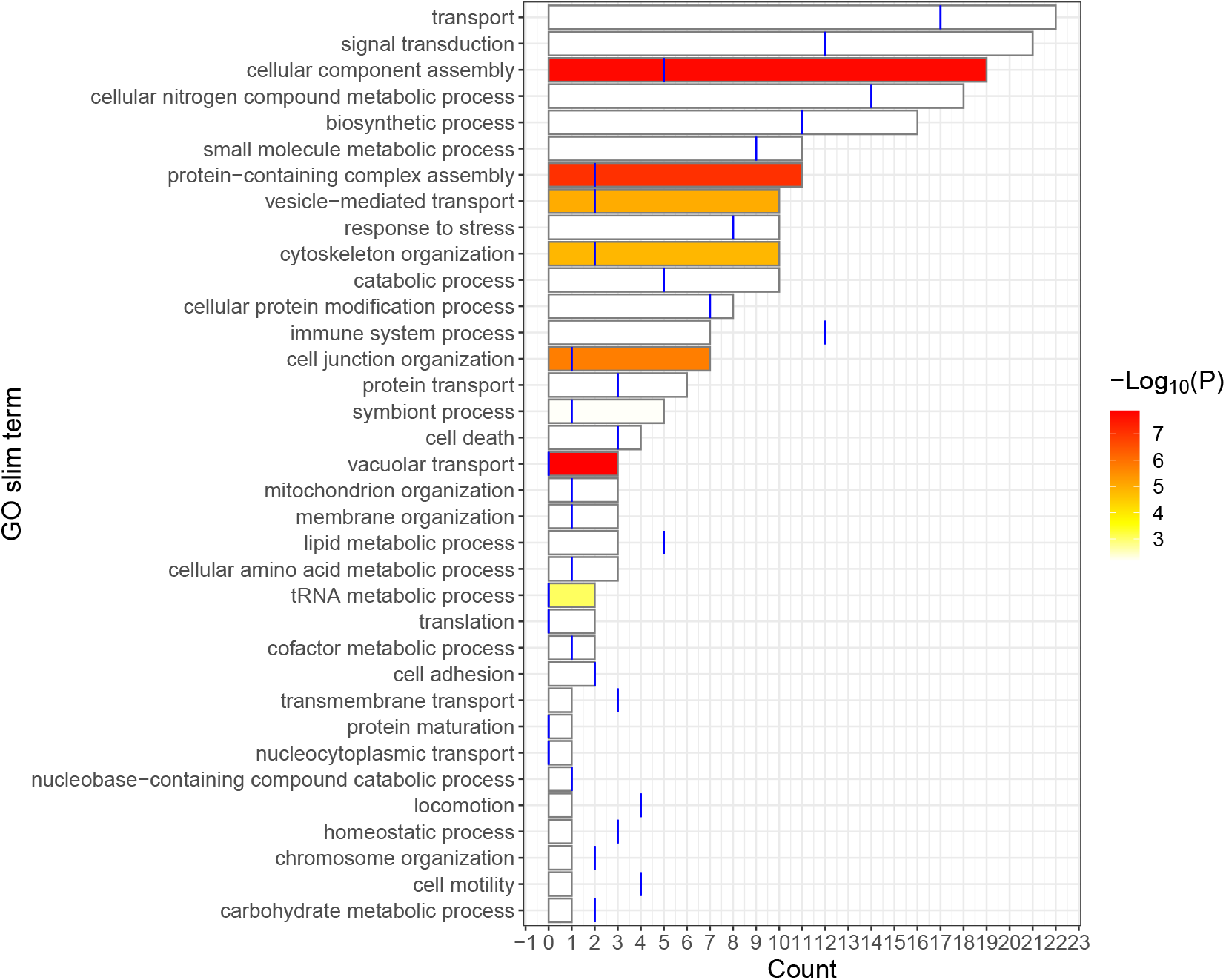
GO slim map and enrichment when the reference population is all datasets combined. The bar graph shows the number of GO terms, out of the 176 terms that overlap in three datasets, that are mapped to the GO slims listed. The expected number of systems mapping to the GO slims is given by the blue line. This was calculated by multiplying the expected proportions of the GO terms tested mapping to the GO slim with the number of unique systems overlapping in at least three datasets (176). The color intensity is the negative log of the p-values from the one-tailed z-tests of proportions. Color is set to white when p-values > 0.05 after adjusting for multiple comparisons with the Benjamini-Hochberg procedure.

**Figure 3–Figure supplement 2.**
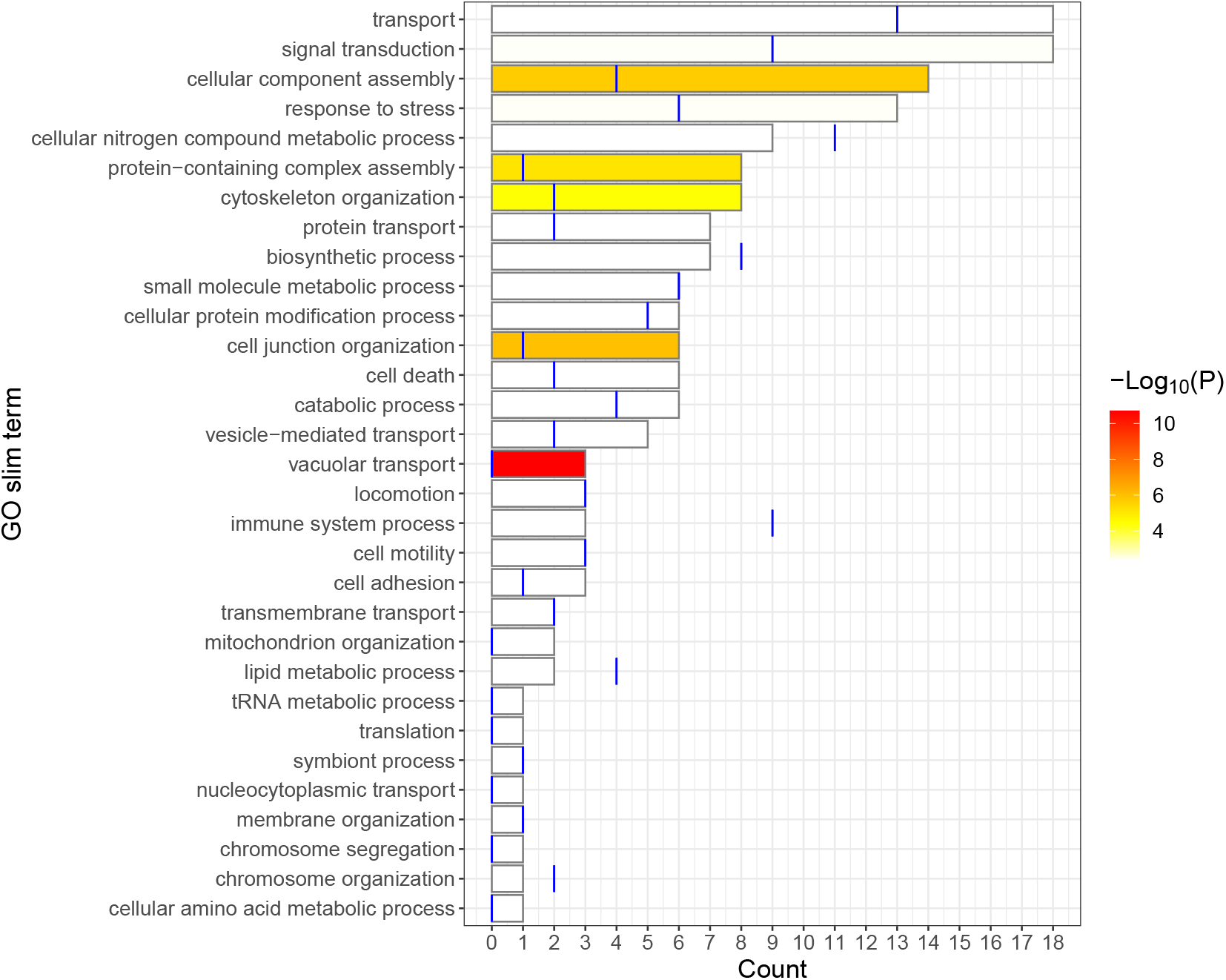
GO slim map and enrichment when the reference population is the Leipzig dataset for all other datasets. The bar graph shows the number of GO terms, out of the 132 terms that overlap in three datasets, that are mapped to the GO slims listed. The expected number of systems mapping to the GO slims is given by the blue line. This was calculated by multiplying the expected proportions of the GO terms tested mapping to the GO slim with the number of unique systems overlapping in at least three datasets (132). The color intensity is the negative log of the p-values from the one-tailed z-tests of proportions. Color is set to white when p-values > 0.05 after adjusting for multiple comparisons with the Benjamini-Hochberg procedure.

**Figure 3–Figure supplement 3.**
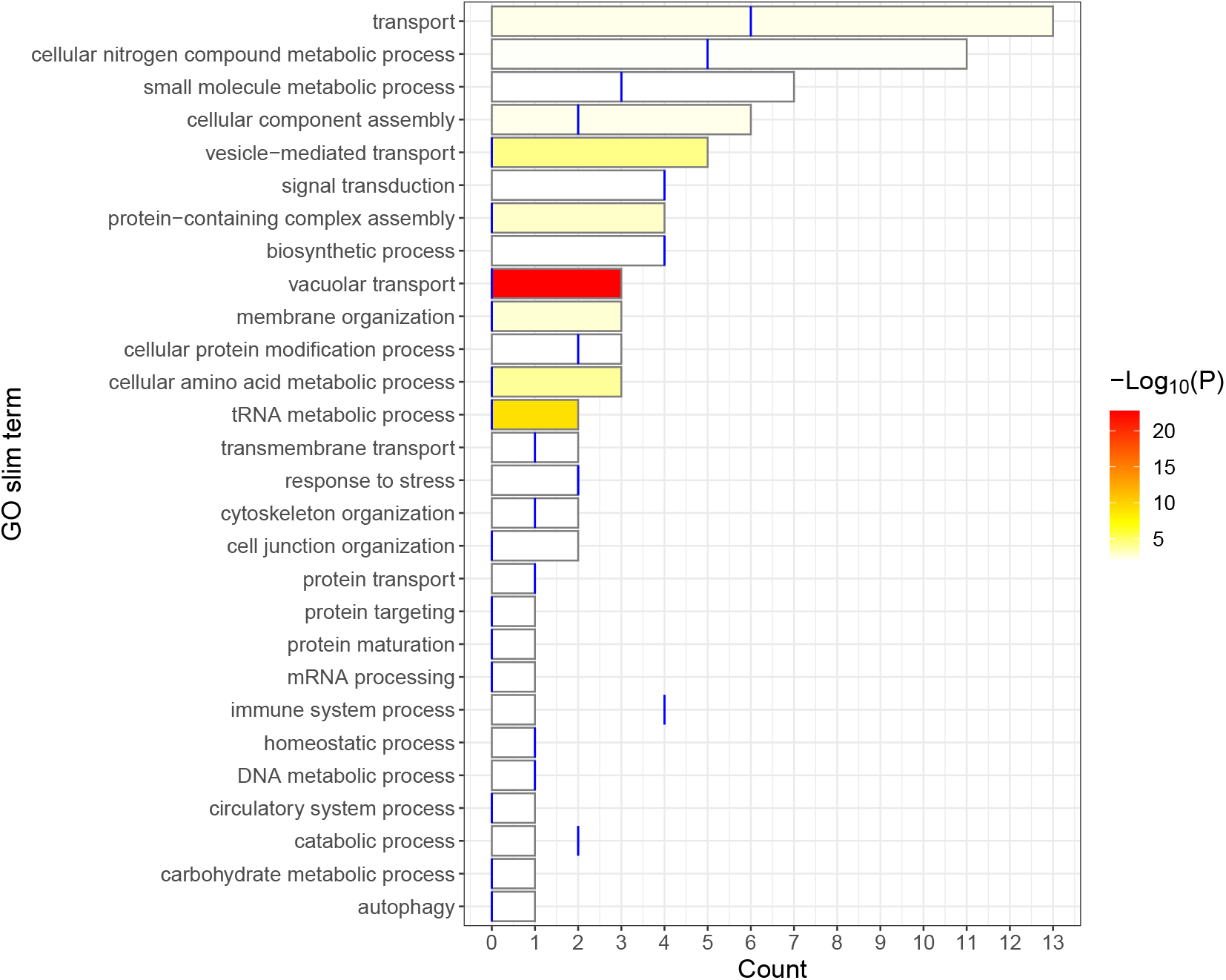
GO slim map and enrichment when the reference population is the AddNeuro dataset for all other datasets. The bar graph shows the number of GO terms, out of the 64 terms that overlap in three datasets, that are mapped to the GO slims listed. The expected number of systems mapping to the GO slims is given by the blue line. This was calculated by multiplying the expected proportions of the GO terms tested mapping to the GO slim with the number of unique systems overlapping in at least three datasets (64). The color intensity is the negative log of the p-values from the one-tailed z-tests of proportions. Color is set to white when p-values > 0.05 after adjusting for multiple comparisons with the Benjamini-Hochberg procedure.

## References

Abbas AR, Baldwin D, Ma Y, Ouyang W, Gurney A, Martin F, Fong S, van Lookeren Campagne M, Godowski P, Williams PM, Chan AC, Clark HF. Immune Response in Silico (IRIS): Immune-Specific Genes Identified from a Compendium of Microarray Expression Data. Genes & Immunity. 2005 Jun; 6(4):319–331. doi: 10.1038/sj.gene.6364173.

Alemu EY, Carl JW, Corrada Bravo H, Hannenhalli S. Determinants of Expression Variability. Nucleic Acids Research. 2014 Apr; 42(6):3503–3514. doi: 10.1093/nar/gkt1364.

Ashburner M, Ball CA, Blake JA, Botstein D, Butler H, Cherry JM, Davis AP, Dolinski K, Dwight SS, Eppig JT, Harris MA, Hill DP, Issel-Tarver L, Kasarskis A, Lewis S, Matese JC, Richardson JE, Ringwald M, Rubin GM, Sherlock G. Gene Ontology: Tool for the Unification of Biology. Nature Genetics. 2000 May; 25(1):25–29. doi: 10.1038/75556.

Barbosa-Morais NL, Dunning MJ, Samarajiwa SA, Darot JFJ, Ritchie ME, Lynch AG, Tavaré S.. A Re-Annotation Pipeline for Illumina BeadArrays: Improving the Interpretation of Gene Expression Data. Nucleic Acids Research. 2010 Jan; 38(3):e17–e17. doi: 10.1093/nar/gkp942.

Belsky DW, Moffitt TE, Cohen AA, Corcoran DL, Levine ME, Prinz JA, Schaefer J, Sugden K, Williams B, Poulton R, Caspi A.. Eleven Telomere, Epigenetic Clock, and Biomarker-Composite Quantifications of Biological Aging: Do They Measure the Same Thing? American Journal of Epidemiology. 2017 Nov; doi: 10.1093/aje/kwx346.

Beutner F, Teupser D, Gielen S, Holdt LM, Scholz M, Boudriot E, Schuler G, Thiery J. Rationale and Design of the Leipzig (LIFE) Heart Study: Phenotyping and Cardiovascular Characteristics of Patients with Coronary Artery Disease. PLoS ONE. 2011 Dec; 6(12):e29070. doi: 10.1371/journal.pone.0029070.

Brinkmeyer-Langford CL, Guan J, Ji G, Cai JJ. Aging Shapes the Population-Mean and -Dispersion of Gene Expression in Human Brains. Frontiers in Aging Neuroscience. 2016 Aug; 8. doi: 10.3389/fnagi.2016.00183.

Chen C, Grennan K, Badner J, Zhang D, Gershon E, Jin L, Liu C. Removing Batch Effects in Analysis of Expression Microarray Data: An Evaluation of Six Batch Adjustment Methods. PLoS ONE. 2011 Feb; 6(2). doi: 10.1371/journal.pone.0017238.

Clough E, Barrett T. The Gene Expression Omnibus Database. Methods in molecular biology (Clifton, NJ). 2016; 1418:93–110. doi: 10.1007/978-1-4939-3578-9_5.

Cohen AA. Complex Systems Dynamics in Aging: New Evidence, Continuing Questions Biogerontology. 2016 Feb; 17(1):205–220. doi: 10.1007/s10522-015-9584-x.

Dunning M, Lynch A, Eldridge M illuminaHumanv4.Db: Illumina HumanHT12v4 Annotation Data (Chip illuminaHumanv4); 2015.

Dunning MJ, Smith ML, Ritchie ME, Tavaré S. Beadarray: R Classes and Methods for Illumina Bead-Based Data. Bioinformatics. 2007; 23(16):2183–4.

Fox RI. Sjögren’s Syndrome. The Lancet. 2005 Jul; 366(9482):321–331. doi: 10.1016/S0140-6736(05)66990-5.

Freytag S, Gagnon-Bartsch J, Speed TP, Bahlo M. Systematic Noise Degrades Gene Co-Expression Signals but Can Be Corrected. BMC Bioinformatics. 2015 Sep; 16(1):1–17. doi: 10.1186/s12859-015-0745-3.

Guan J, Cai JJ, Ji G, Sham PC Commonality in Dysregulated Expression of Gene Sets in Cortical Brains of Individuals with Autism, Schizophrenia, and Bipolar Disorder. Translational Psychiatry. 2019 Dec; 9(1):152. doi: 10.1038/s41398-019-0488-4.

Guan J, Yang E, Yang J, Zeng Y, Ji G, Cai JJ. Exploiting Aberrant mRNA Expression in Autism for Gene Discovery and Diagnosis. Human Genetics. 2016 Jul; 135(7):797–811. doi: 10.1007/s00439-016-1673-7.

Huang G, Osorio D, Guan J, Ji G, Cai JJ. Overdispersed Gene Expression in Schizophrenia. npj Schizophrenia. 2020 Dec; 6(1):9. doi: 10.1038/s41537-020-0097-5.

Ikram MA, Brusselle GGO, Murad SD, van Duijn CM, Franco OH, Goedegebure A, Klaver CCW, Nijsten TEC, Peeters RP, Stricker BH, Tiemeier H, Uitterlinden AG, Vernooij MW, Hofman A The Rotterdam Study: 2018 Update on Objectives, Design and Main Results. European Journal of Epidemiology. 2017 Sep; 32(9):807–850. doi: 10.1007/s10654-017-0321-4.

Işıldak U, Somel M, Thornton JM,. Dönertaş HM. Temporal Changes in the Gene Expression Heterogeneity during Brain Development and Aging. Scientific Reports. 2020 Dec; 10(1):4080. doi: 10.1038/s41598-020-60998-0.

Johnson WE, Li C, Rabinovic A. Adjusting Batch Effects in Microarray Expression Data Using Empirical Bayes Methods. Biostatistics. 2007 Jan; 8(1):118–127. doi: 10.1093/biostatistics/kxj037.

Kauffman SA, et al. The Origins of Order: Self-Organization and Selection in Evolution. Oxford University Press, USA; 1993.

Kedlian VR, Donertas HM, Thornton JM. The Widespread Increase in Inter-Individual Variability of Gene Expression in the Human Brain with Age. Aging. 2019 Apr; 11(8):2253–2280. doi: 10.18632/aging.101912.

Kimmel JC, Penland L, Rubinstein ND, Hendrickson DG, Kelley DR, Rosenthal AZ. Murine Single-Cell RNA-Seq Reveals Cell-Identity- and Tissue-Specific Trajectories of Aging. Genome Research. 2019 Dec; 29(12):2088–2103. doi: 10.1101/gr.253880.119.

Kirkwood TBL. Understanding the Odd Science of Aging. Cell. 2005 Feb; 120(4):437–447. doi: 10.1016/j.cell.2005.01.027.

Leek JT, Johnson WE, Parker HS, Fertig EJ, Jaffe AE, Storey JD, Zhang Y, Torres LC Sva: Surrogate Variable Analysis; 2019.

Li Q, Wang S, Milot E, Bergeron P, Ferrucci L, Fried LP, Cohen AA. Homeostatic Dysregulation Proceeds in Parallel in Multiple Physiological Systems. Aging Cell. 2015 Dec; 14(6):1103–1112. doi: 10.1111/acel.12402.

Li Z, Wright FA, Royland J. Age-Dependent Variability in Gene Expression in Male Fischer 344 Rat Retina. Toxicological Sciences. 2009 Jan; 107(1):281–292. doi: 10.1093/toxsci/kfn215.

López-Otí C, Blasco MA, Partridge L, Serrano M, Kroemer G. The Hallmarks of Aging. Cell. 2013 Jun; 153(6):1194–1217. doi: 10.1016/j.cell.2013.05.039.

Mahalanobis PC. On the Generalized Distance in Statistics. Proceedings of the National Institute of Sciences (Calcutta). 1936; 2:49–55.

Milot E, Morissette-Thomas V, Li Q, Fried LP, Ferrucci L, Cohen AA. Trajectories of Physiological Dysregulation Predicts Mortality and Health Outcomes in a Consistent Manner across Three Populations. Mechanisms of ageing and development. 2014; 141-142:56–63. doi: 10.1016/j.mad.2014.10.001.

Müller C, Schillert A, Röthemeier C, Trégouët DA, Proust C, Binder H, Pfeiffer N, Beutel M, Lackner KJ, Schnabel RB, Tiret L, Wild PS, Blankenberg S, Zeller T, Ziegler A. Removing Batch Effects from Longitudinal Gene Expression - Quantile Normalization Plus ComBat as Best Approach for Microarray Transcriptome Data. PLoS ONE. 2016 Jun; 11(6):1–23. doi: 10.1371/journal.pone.0156594.

Peters MJ, Pilling LC, Schurmann C, Conneely KN, Powell J, Reinmaa E, Sutphin GL, Zhernakova A, Schramm K, Wilson YA, Kobes S, Tukiainen T, Ramos YF, Göring HHH, Fornage M, Liu Y, Gharib SA, Stranger BE, De Jager PL, Aviv A, et al. The Transcriptional Landscape of Age in Human Peripheral Blood. Nature Communications. 2015 Dec; 6(1):8570. doi: 10.1038/ncomms9570.

R Core Team. R: A Language and Environment for Statistical Computing. Vienna, Austria; 2019.

Reimers M, Carey VJ. Bioconductor: An Open Source Framework for Bioinformatics and Computational Biology. Methods in Enzymology. 2006; 411:119–134. doi: 10.1016/S0076-6879(06)11008-3.

Ritchie ME, Phipson B, Wu D, Hu Y, Law CW, Shi W, Smyth GK. Limma Powers Differential Expression Analyses for RNA-Sequencing and Microarray Studies. Nucleic Acids Research. 2015; 43(7):e47. doi: 10.1093/nar/gkv007.

Sarkans U, Parkinson H, Lara GG, Oezcimen A, Sharma A, Abeygunawardena N, Contrino S, Holloway E, Rocca-Serra P, Mukherjee G, Shojatalab M, Kapushesky M, Sansone SA, Farne A, Rayner T, Brazma A. The ArrayExpress Gene Expression Database: A Software Engineering and Implementation Perspective. Bioinformatics. 2005 Apr; 21(8):1495–1501. doi: 10.1093/bioinformatics/bti157.

Schissler AG, Gardeux V, Li Q, Achour I, Li H, Piegorsch WW, Lussier YA. Dynamic Changes of RNA-Sequencing Expression for Precision Medicine: N-of-1-Pathways Mahalanobis Distance within Pathways of Single Subjects Predicts Breast Cancer Survival. Bioinformatics. 2015 Jun; 31(12):i293–i302. doi: 10.1093/bioinformatics/btv253.

Sood S, Gallagher IJ, Lunnon K, Rullman E, Keohane A, Crossland H, Phillips BE, Cederholm T, Jensen T, van Loon LJ, Lannfelt L, Kraus WE, Atherton PJ, Howard R, Gustafsson T, Hodges A, Timmons JA. A Novel Multi-Tissue RNA Diagnostic of Healthy Ageing Relates to Cognitive Health Status. Genome Biology. 2015 Dec; 16(1):185. doi: 10.1186/s13059-015-0750-x.

Stein PK, Barzilay JI, Chaves PHM, Domitrovich PP, Gottdiener JS. Heart Rate Variability and Its Changes over 5 Years in Older Adults. Age and Ageing. 2009 Mar; 38(2):212–218. doi: 10.1093/ageing/afn292.

Tarn JR, Howard-Tripp N, Lendrem DW, Mariette X, Saraux A, Devauchelle-Pensec V, Seror R, Skelton A J, James K, McMeekin P, Al-Ali S, Hackett KL, Lendrem BC, Hargreaves B, Casement J, Mitchell S, Bowman SJ, Price E, Pease CT, Emery P, et al. Symptom-Based Stratification of Patients with Primary Sjögren’s Syndrome: Multi-Dimensional Characterisation of International Observational Cohorts and Reanalyses of Randomised Clinical Trials. The Lancet Rheumatology. 2019 Oct; 1(2):e85–e94. doi: 10.1016/S2665-9913(19)30042-6.

Valiathan R, Ashman M, Asthana D. Effects of Ageing on the Immune System: Infants to Elderly. Scandinavian Journal of Immunology. 2016; 83(4):255–266. doi: 10.1111/sji.12413.

Viñuela A, Brown AA, Buil A, Tsai PC, Davies MN, Bell JT, Dermitzakis ET, Spector TD, Small KS. Age-Dependent Changes in Mean and Variance of Gene Expression across Tissues in a Twin Cohort. Human Molecular Genetics. 2018 Feb; 27(4):732–741. doi: 10.1093/hmg/ddx424.

Wang M, Zhao Y, Zhang B SuperExactTest: Exact Test and Visualization of Multi-Set Intersections; 2019.

Wei T, Simko V R Package “Corrplot”: Visualization of a Correlation Matrix; 2017.

Wickham H, Averick M, Bryan J, Chang W, McGowan L, François R, Grolemund G, Hayes A, Henry L, Hester J, Kuhn M, Pedersen T, Miller E, Bache S, Müller K, Ooms J, Robinson D, Seidel D, Spinu V, Takahashi K, et al. Welcome to the Tidyverse. Journal of Open Source Software. 2019; 4(43):1686. doi: 10.21105/joss.01686.

Wingo AP, Gibson G. Blood Gene Expression Profiles Suggest Altered Immune Function Associated with Symptoms of Generalized Anxiety Disorder. Brain, behavior, and immunity. 2015 Jan; 43:184–191. doi: 10.1016/j.bbi.2014.09.016.

Zeng Y, Wang G, Yang E, Ji G, Brinkmeyer-Langford CL, Cai JJ. Aberrant Gene Expression in Humans. PLOS Genetics. 2015 Jan; 11(1):e1004942. doi: 10.1371/journal.pgen.1004942.

